# Tunneling nanotubes between neuronal and microglial cells allow bi-directional transfer of *α*-Synuclein and mitochondria

**DOI:** 10.1101/2022.12.13.519450

**Authors:** Ranabir Chakraborty, Chiara Zurzolo

## Abstract

Tunneling Nanotubes (TNTs) facilitate contact-mediated intercellular communication over long distances. Material transfer via TNTs can range from ions and intracellular organelles to protein aggregates and pathogens. Prion-like toxic protein aggregates accumulating in several disorders, such as Alzheimer’s, Parkinson’s, and Huntington’s diseases have been shown to spread via TNTs not only between neurons, but also between neurons-astrocytes, and neurons-pericytes, indicating the importance of TNTs in mediating neuron-glia interactions. TNT-like structures were also reported between microglia, however their roles in neuron-microglia interaction remain elusive. In this work, we quantitatively characterise microglial TNTs and their cytoskeletal composition, and demonstrate that TNTs form between human neuronal and microglial cells. We show that *α*-Synuclein (*α*-Syn) aggregates increase the global TNT-mediated connectivity between cells, along with the number of TNT connections per cell pair. Homotypic TNTs formed between microglial cells, and heterotypic TNTs between neuronal and microglial cells are furthermore shown to be functional, allowing movement of both *α*-Syn and mitochondria. Quantitative analysis shows that *α*-Syn aggregates are transferred predominantly from neuronal to microglial cells, possibly as a mechanism to relieve the burden of accumulated aggregates. By contrast, microglia transfer mitochondria preferably to *α*-Syn burdened neuronal cells over the healthy ones, likely as a potential rescue mechanism. Besides describing novel TNT-mediated communication between neurons and microglia, this work allows us to better understand the cellular mechanisms of spreading of neurodegenerative diseases, shedding light on the role of microglia.

## Introduction

Accumulation of neurotoxic protein aggregates in neurodegenerative diseases (NDDs) such as Alzheimer’s disease (AD), Parkinson’s disease (PD), Huntington’s disease (HD), eventually lead to the death of burdened neurons that are unable to dilute the aggregates because of their post-mitotic nature, as well as because of the impairment of the degradative pathways (1,2). Although cells have different quality control mechanisms in place to clear misfolded proteins, such as the ubiquitin-proteasome system (UPS), and autophagy, continuous generation of aggregates renders these processes ineffective in maintaining proteostasis (2,3). Besides the role of intrinsic protein accumulation, neuronal death in NDDs is also regulated in a non-cell autonomous manner by glial cells. Both astrocytes, the major glial cell type, and microglia, the tissue-resident macrophages of the brain respond to neurodegenerating conditions by altering their cellular properties from a homeostatic to a reactive phenotype, a phenomenon characteristic of neuroinflammation (4). If extended beyond its initial stages of brain repair and neuroprotection, neuroinflammation can result in progression of neurodegeneration (5).

Alongside astrocytes, microglia are major responders to impaired neuronal proteostasis. Presence of protein aggregates drives microglia towards a reactive phenotype, that subsequently initiates various direct and indirect routes leading to neuronal death (6). In cases of PD, degeneration of dopaminergic neurons begin at the substantia nigra pars compacta (SNpc). In later stages, the pathology spreads and *α*-Syn deposition has been observed in different other regions of the brain, implicating the propagative nature of these aggregates in a “prion-like” mechanism (7–9). A mechanism of disease propagation is via the release of *α*-Syn aggregates by neurons, which eventually binds to, and is engulfed by, surrounding neurons (10–13). Besides neurons, neighbouring astrocytes and microglia express cell-surface receptors that recognise *α*-Syn as a ligand, such as toll-like receptor 2 (TLR2), Fyn kinase and scavenger receptor CD36, and FcγRIIB, that drives an NLRP3 inflammasome-driven proinflammatory signaling cascade (14–18). As a consequence, microglia in different regions eventually acquire a reactive phenotype. Furthermore, upon binding of neuron-released *α*-Syn to TLR4, microglia have been reported to upregulate the expression of p62/SQSTM in an NF*k*B-dependent manner that eventually leads to the autophagic clearance of the aggregates (19).

In addition to secretion-based mechanisms, recent *in vitro and ex vivo* evidence support the involvement of tunneling nanotubes (TNTs) in spreading of neurodegenerative pathologies. First described in 2004, TNTs are thin, membrane-enclosed, actin-rich protrusions that connect cells over long distances (20). Functionally, TNTs are able to transfer cargoes of different kinds between the connected cells, like Ca^2+^ signals, messenger- and micro-RNAs, organelles such as lysosomes and mitochondria, pathogens, apoptotic signals, and NDD-protein aggregates (21,22). Amyloidogenic proteins such as prions (23), amyloid *β*(24), tau (25), mHTT (26), and *α*-Syn (27) have been reported to use TNTs as a route of spreading from one cell to another in co-cultures *in vitro* and in brain slices. Interestingly, *α*-Syn aggregates closely associate with lysosomes, to move to an uninfected neuronal cell and initiate aggregate seeding (28). *α*-Syn fibrils can also propagate between neurons and astrocytes (29,30), as well as neurons and pericytes (31) via tunneling nanotubes (TNTs). Recently, an elegant work using mouse primary microglia, human monocyte-derived microglia like cells, and post-mortem human brain samples, provided qualitative data showing formation of TNTs allowing the spreading of *α*-Syn aggregates between microglial cells and leading to their eventual degradation (32). However, to understand the role of microglia in the progression of NDDs, it is of capital importance to investigate whether TNTs form between neurons and microglia and to what extent do these aggregates move between neurons and microglia and use TNTs to do so.

Using both quantitative and qualitative live imaging approaches, we report here for the first time the presence of functional TNTs between neuronal and microglial cells that allow movement of mitochondria and *α*-Syn aggregates between these cells both uni- and bi-directionally. We quantitatively assess the extent of such transfers, revealing a bias of *α*-Syn transfer from neuronal cells to microglia, and mitochondrial movement from microglia preferably to unhealthy neuronal cells. Our results uncover TNTs as a major route of neuron-microglia interactions in normal and neurodegenerating contexts and support a rescue function for microglia, thereby advancing our understanding neuron-glia interactions.

## Materials and Methods

### Cell culture and α-Syn treatment

Human neuroblastoma cell line SH-SY5Y were grown in RPMI1640 media (Euroclone), supplemented with 10 % FCS and 1 % Penicillin-Streptomycin (Pen-Strep). Human microglial clone 3 (HMC3) cell line, a kind gift from Dr Aleksandra Deczkowska, Institut Pasteur, were grown in DMEM media (Sigma-Aldrich), supplemented with 10 % FCS and 1 % Penicillin-Streptomycin (Pen-Strep). Cells were maintained in a humidified incubator at 37°C, passaged at 80-90 % confluency. Cells were seeded at 1:5 ratio for maintenance in culture, and counted before seeding for particular experiments. *α*-Syn fibrils were a kind gift from Dr Takashi Nonaka and Dr Masato Hasegawa (Dementia Research Project, Tokyo Metropolitan Institute of Medical Science, Tokyo, Japan) and were prepared as originally described (33).

For treatment with *α*-Syn, cells were seeded in uncoated 6-well plates (Falcon) at a density of 400,000 cells/well, and grown for 24 hours (h). *α*-Syn fibrils were prepared as previously described (28). Briefly, fluorophore-conjugated fibrils were diluted in growth medium (500 nM) and sonicated (BioBlock Scientific, Vibra Cell 75041) for 5 minutes (80% amplitude, pulse-on: 5 seconds, off: 2 seconds). Sonicated fibrils were then added to the cells for 16 h of incubation. After 16 h, cells were washed thrice with 1:3 trypsin solution diluted in 1X PBS, and processed further for subsequent experiments.

### Labelling TNTs

Owing to the fragile nature of TNTs, cells were handled with utmost care to minimize physical stress. After cells reached sub-confluency, a two-step fixation protocol was used to preserve TNTs as described previously (34). Briefly, cells were fixed initially with a fixative containing glutaraldehyde (GA) (0.05 % GA, 2 % paraformaldehyde (PFA), 0.2 M HEPES buffer in 1X PBS) for 15 minutes at room temperature (RT), followed by a second fixation without GA for 15 minutes at RT (4 % PFA, 0.2 M HEPES buffer in 1X PBS). Following this, cells were washed thrice with 1X PBS, and stained for plasma membrane with 3.33 *µ*g/mL wheat germ agglutinin (WGA, Life Technologies, Thermo Fisher Scientific), and/or F-Actin with Phalloidin (1:250 v/v, diluted in 1X PBS, Invitrogen, Thermo Fisher Scientific) for 15 minutes at RT in dark. For F-Actin staining in Fig. 3, CellMask deep red Actin tracking stain (1:1000 dilution, Invitrogen) was used.

### Immunocytochemistry

Cells were grown to sub-confluency and rinsed once with freshly-prepared and warmed cytoskeleton buffer (60 mM PIPES buffer, pH 6.9; 25 mM HEPES, 10 mM EGTA and 2 mM MgCl_2_ in milli-Q water). Cells were then fixed with 0.05 % GA and 4 % PFA diluted in cytoskeleton buffer for 20 minutes at 37°C. PFA was quenched with 50 mM NH_4_Cl for 15 minutes at RT, followed by three washes with 1X PBS. Cells were then permeabilized with 0.1 % Triton X-100 in 1X PBS (0.1 % PBSTx) for 3 minutes, followed by blocking with 2 % bovine serum albumin (BSA, Sigma Aldrich). Primary antibody against *α*-Tubulin was diluted (1:500, Sigma Aldrich) in 2% BSA and cells were incubated overnight at 4°C. The next day, cells were washed thrice with 0.1 % PBSTx, and incubated with AlexaFluor488-conjugated secondary antibody diluted in 2 % BSA (1:500, Invitrogen, Thermo Fisher Scientific) against the primary antibody for 40 minutes at RT. Cells were then washed thrice with 0.1 % PBSTx and incubated with AlexaFluor568-conjugated Phalloidin (1:250 v/v, diluted in 1X PBS) for 15 minutes at RT in dark. Cells were washed thrice with 1X PBS, incubated with AlexaFluor647-conjugated WGA (3.33 *µ*g/mL diluted in 1X PBS) for 15 minutes at RT, washed thrice with 1X PBS before mounting with Aqua-Poly/Mount (Polysciences).

### Transfer assay of α-Syn

To assess for the movement of *α*-Syn between neuronal and microglial cells, a co-culture strategy was adopted. Assessing for transfer from neuronal cells to microglia (N→M), SH-SY5Y were loaded with *α*-Syn fibrils (donor cells) and co-cultured in a 1:1 ratio with HMC3 (acceptor cells) on coverslips in 24-well plates. In another condition to check for transfer in the opposite direction, microglia were the donor cells while neuronal cells were acceptors (M→N). Cells were grown for 24 h, fixed to preserve TNTs (as mentioned previously), stained for membrane, and mounted on slides.

To negate the possibility of transfer in a secretion-dependent manner, conditioned medium from donor cells grown in mono-culture were added to acceptor cells for 24 h, followed by fixation and mounting.

### Transfer assay of mitochondria

Transfer of mitochondria was assessed from microglia to neuronal cells with or without *α*-Syn (referred as M→N (+*α*-Syn) and M→N (WT) respectively). Microglial mitochondria were labelled with MitoTracker Red CMXRos (500 nM, Invitrogen, Thermo Fisher Scientific) diluted in growth medium for 30 minutes at 37°C before seeding for co-culture. Neuronal cells (WT or +*α*-Syn; acceptor cells) were co-cultured in a 1:1 ratio with microglia (donor cells) and grown for 24 h before fixation as mentioned previously.

For secretion control, conditioned medium from mitochondria-labelled microglia grown in mono-culture was added to acceptor cells (WT and +*α*-Syn) for 24 h before fixation and mounting.

### Fixed-cell imaging

Images were acquired using Zeiss LSM900 confocal microscope, 40X/1.3 NA oil objective and 0.8x zoom. All images were acquired with the “confocal” setting for optimal pixel sampling of the regions of interests. Optical slices were designated at a constant interval of 0.45 *µ*m.

### Live-cell imaging

For live cell imaging, cells were grown in 35 mm glass-bottom dishes (Ibidi) for 24 h. Cells were loaded with CellMask Actin tracking stain (diluted in growth media to a working concentration of 1X from a 1000X stock solution) for 30 minutes at 37°C. Cells were washed thrice with growth media before imaging. Time-lapse images were acquired using Nikon Eclipse Ti2 spinning disk microscope, 60X/1.4 NA oil objective every minute for 30 minutes (30 frames). Optical sections were set at optimum for acquisition.

### Image analysis and statistics

All time-fixed and time-lapse images were analysed using FIJI image processing software. For analysing the proportion of TNT-connected cells, images were analysed using ICY software (https://icy.bioimageanalysis.org/). Movies were created at a rate of 2 frames per second. Graphs were plotted in GraphPad Prism 7.0, and appropriate statistical tests were implemented, as mentioned in the figure legends. Statistical significance was calculated with an *α*-value of 0.05. To determine the effect sizes and calculate Cohen’s d, Bland-Altmann plots were plotted with the estimation statistics online tool (www.estimationstats.com; (35)).

## Results

### Tunneling nanotubes connect HMC3 microglia

A recent report provides evidence of TNT-like structures connecting murine primary microglia and human monocyte-derived microglia like cells (32). Because no specific biomarker exists for these structures, the categorisation of cellular protrusions as TNTs requires several stringent criteria to be considered (34,36). Accordingly, our morphological identification of TNTs refers to membranous intercellular connections with a length of 10 *µ*m and more (to differentiate them from filopodia) (37,38) that hover above the substratum, and contain F-Actin. Using HMC3 microglial cell line, stained for membrane and F-Actin (Fig. 1A), we show that under normal growing conditions, 39.82 % of HMC3 cells were connected by TNTs (Fig. 1B), with majority of their lengths being in a range of 10-20 *µ*m (Fig. 1B, C) supporting the qualitative evidence of the existence of TNTs between microglia both *in vitro*, and in post-mortem human brain samples (32).

**Fig. 1:**
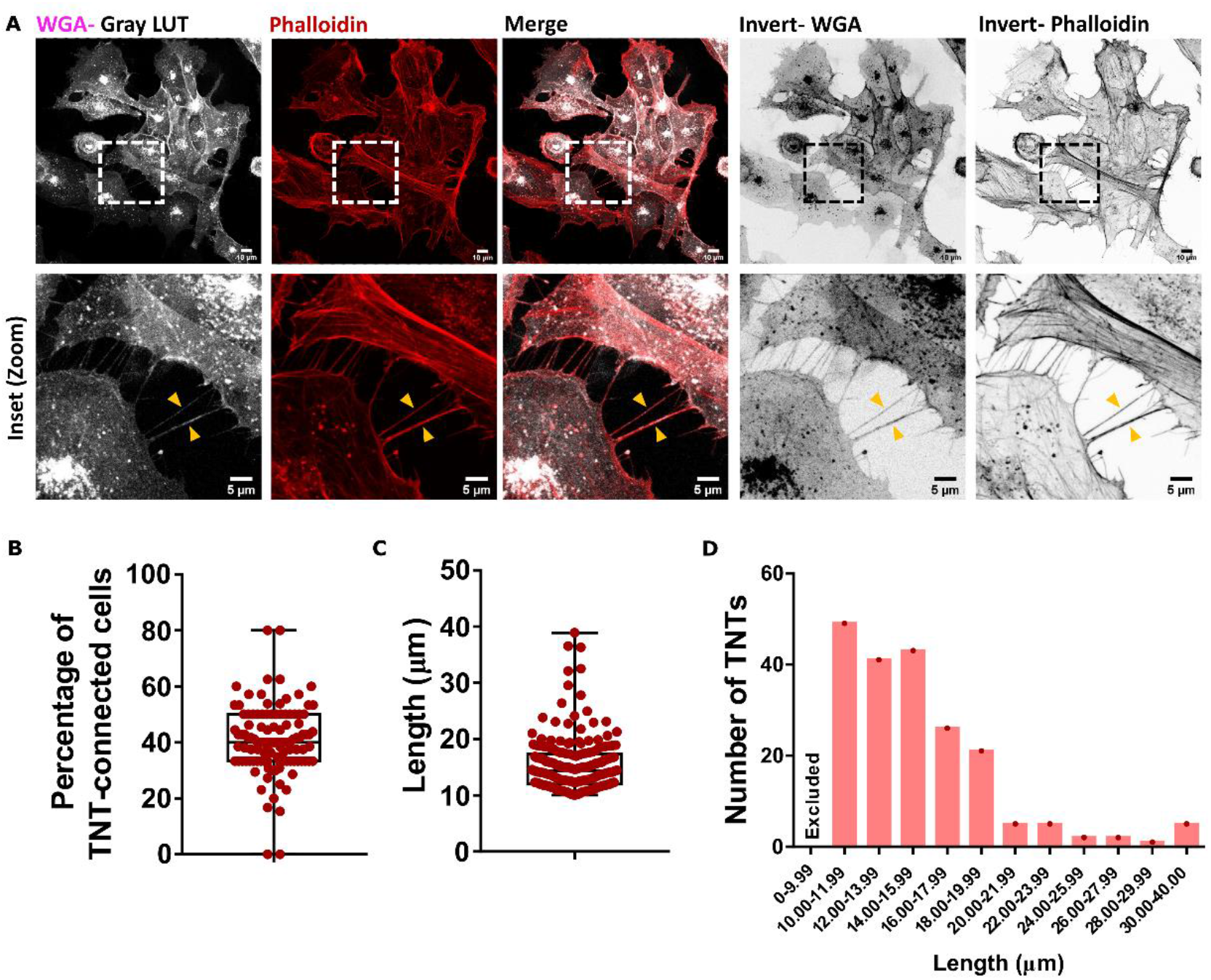
Tunneling nanotubes are present between microglia. (A) HMC3 microglia stained for membrane with AlexaFluor647-conjugated WGA (pseudo-coloured gray) and F-Actin with Rhodamine phalloidin (red). Bottom panel represents the zoomed region depicted with boxes in the upper panel. Yellow arrowheads point towards TNTs. (B) Proportion of TNT-connected cells under normal growing conditions (N=3, n=111 regions of interests). (C) Lengths of the TNTs observed. (D) Distribution of TNTs based on their lengths. Structures less than 10 *µ*m were excluded from our analyses. N=3, n=200 TNTs for (C) and (D).

### Microglial TNTs contain F-Actin alone, or together with α-Tubulin

Although ‘thin’ or ‘canonical’ TNTs have been described to be rich in F-Actin filaments for a variety of cell types, such as SH-SY5Y neuroblastoma cells (39), the presence of microtubule along with F-Actin have been reported in ‘thick’ TNTs, characteristic of cells of myeloid origin (40,41). Of interest, quantitative assessment of the cytoskeletal composition of TNTs, showed that, differently from myeloid cells, most (70.74 %) of the microglia TNTs contained only F-Actin (Fig. 2A, E). However, 29.26 % of the TNTs contained microtubule along with F-Actin, albeit to different extents. In 25.56 % of the TNTs, microtubule presence was detected at a very low level, visualised evidently only upon digital overexposure (Fig. 1D, E). In 2.22 % of the cases, TNTs contained microtubule only up to a segment of the structure (Fig. 1C, E), whereas 1.48 % of the TNTs contained microtubule throughout the length (Fig. 1 B, E).

**Fig. 2:**
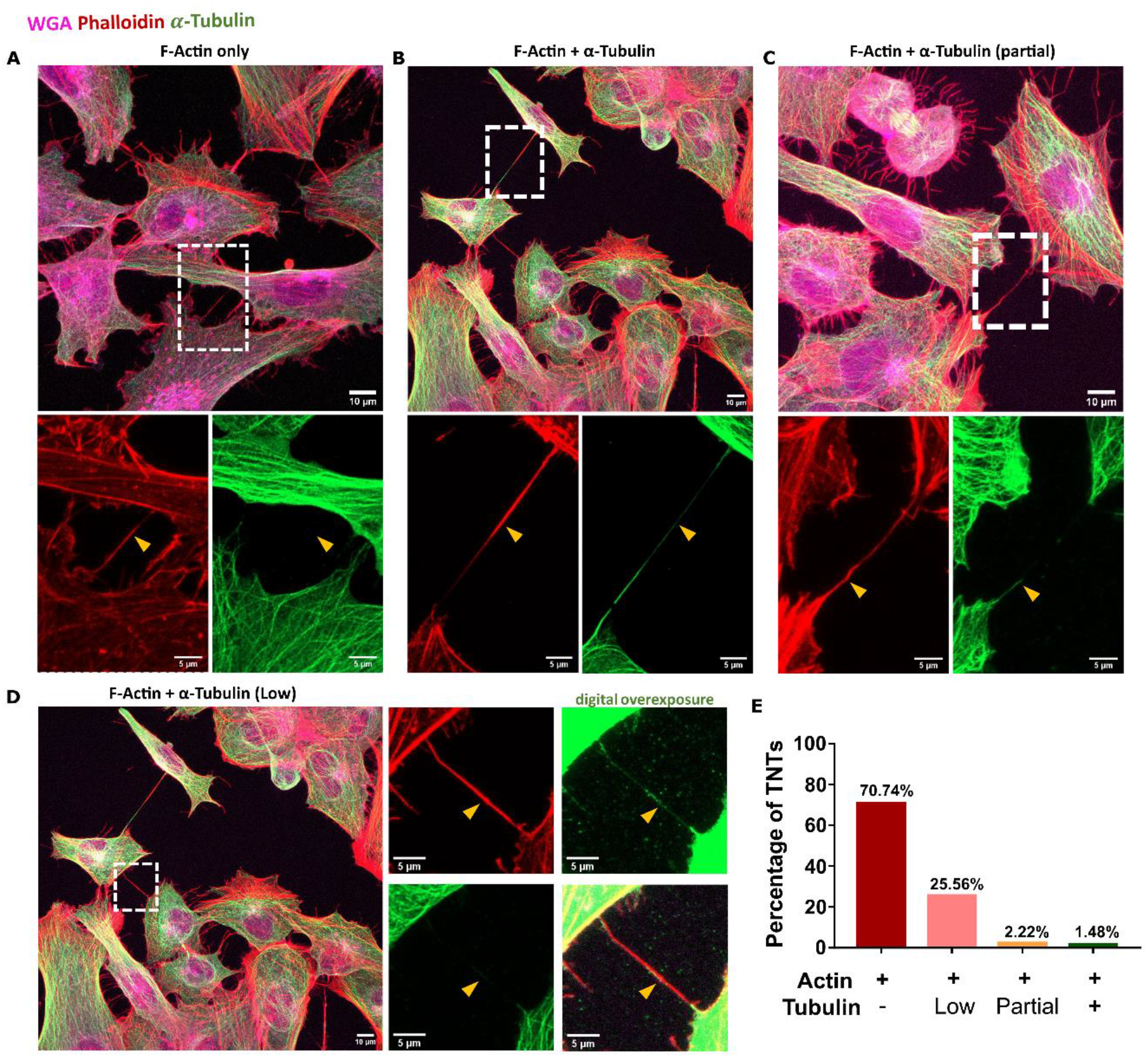
Cytoskeletal composition of microglial TNTs. (A-D) Immunostaining for *α*-Tubulin (green) and subsequent staining for F-Actin with Rhodamine phalloidin (red) and membrane with AlexaFluor647-conjugated WGA (magenta). (A) TNTs containing only F-Actin. (B) TNTs containing both F-Actin and *α*-Tubulin. (C) TNTs partially containing *α*-Tubulin. (D) TNTs containing low level of *α*-Tubulin. (E) Proportion of TNTs with diverse cytoskeletal compositions. N=3, n=270 TNTs.

**Fig. 3:**
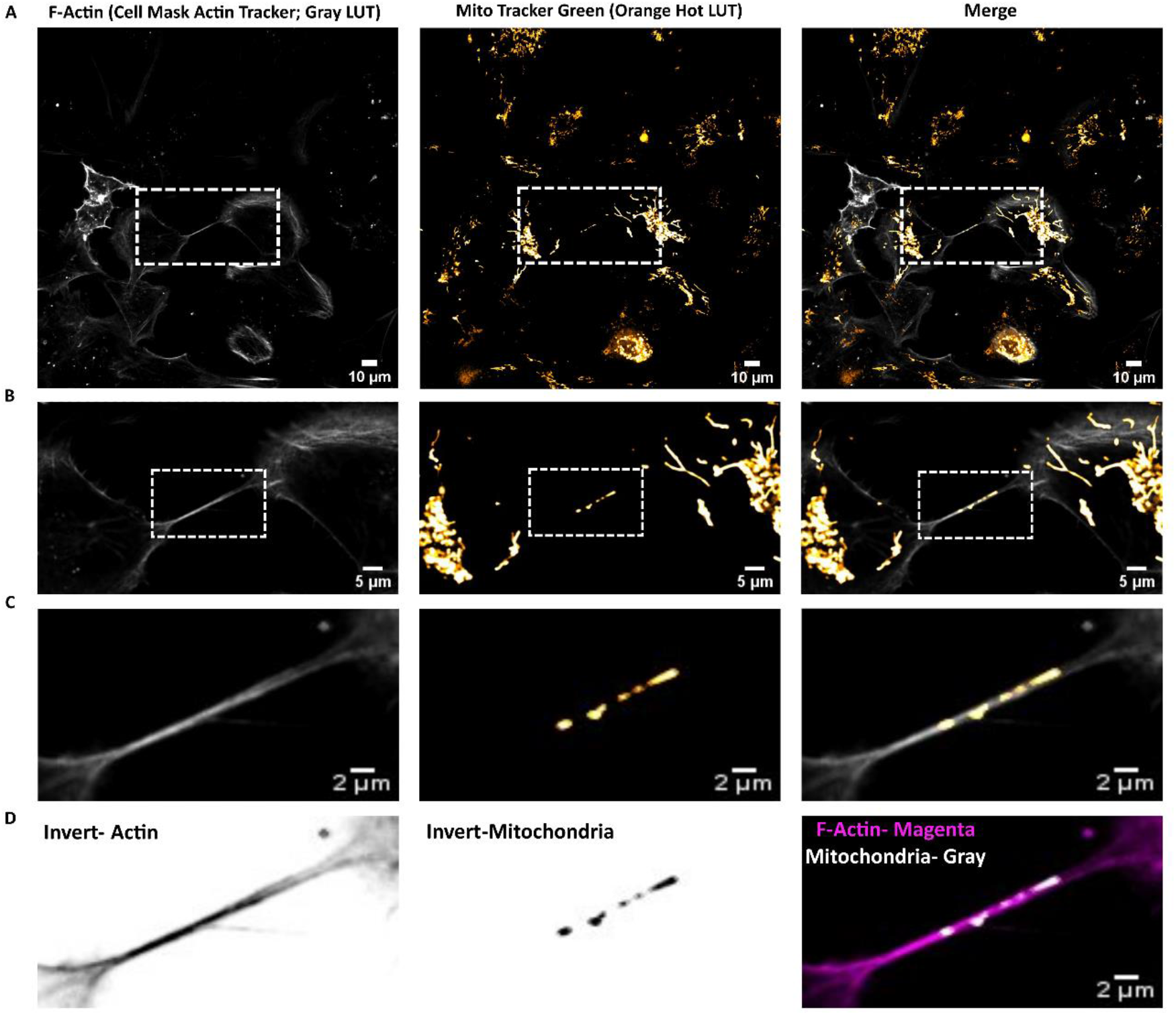
Functional TNTs are formed between microglia. (A) Single, middle stack images of microglial cells stained for F-Actin with CellMask deep red Actin tracking stain (pseudo-coloured gray; left panels) and mitochondria with MitoTracker green FM (pseudo-coloured orange, middle panels). (B) Zoomed images from the ROIs boxed in (A). (C-D) Zoomed images from the ROIs boxed in (B). Image snapshots were acquired using Nikon Eclispe Ti2 spinning disk microscope.

### TNTs formed by microglia contain mitochondria

Despite these morphological characteristics, an essential requirement for intercellular connections to be classified as TNTs is their ability to transfer cargoes between the cells. A major intracellular component commonly reported to be transferred via TNTs between different cell types and experimental conditions is mitochondria (reviewed in (22)). Towards this end, the presence of mitochondrial particles inside of F-Actin containing TNTs connecting two microglia (Fig. 3A-D) supported the functional nature of these TNTs, which was further assessed by live imaging (see below, Supplementary movie 3).

### Functional TNTs are formed between neuronal cells and microglia

Although TNTs have been reported to form between neurons, and between microglia, in homotypic cultures, their presence between neurons and microglia has not been assessed. Using a co-culture strategy between human neuronal SH-SY5Y cells and HMC3 microglial cells(Fig. 4A), we could observe TNTs connecting the two cell populations. To assess the functionality of these TNTs, we loaded SH-SY5Y cells with *α*-Syn (donor cells, Fig. S1A, upper panels) and measured quantitatively the transfer of these aggregates to microglia (acceptor cells, Fig. S1A, upper panels) (Fig. 4C, F, G). We found that 57.85 % of acceptor cells were positive for *α*-Syn puncta (N→M transfer) (Fig. 4F). However, when microglia were the donor cells and neuronal cells were acceptors (Fig. S1A lower panels) (M→N transfer), the extent of transfer was limited to 10.02 %, lesser by 5.87 folds (Fig 4D, F, G, H). Orthogonal projection (Fig. 4C) and 3-D reconstruction of the TNT connecting donor neuronal cells with acceptor microglia (Fig. 4E) demonstrate the presence of *α*-Syn within the TNT. With a high value of Cohen’s d (Fig. 4K), the effect size of the difference was substantial. Additionally, the number of *α*-Syn puncta per acceptor cell were higher in microglia as opposed to neuronal cells (Fig. 4G, I, L), with a 3.04-folds increase in acceptor microglial cells compared to acceptor neuronal cells (Fig. 4I).

**Fig. 4:**
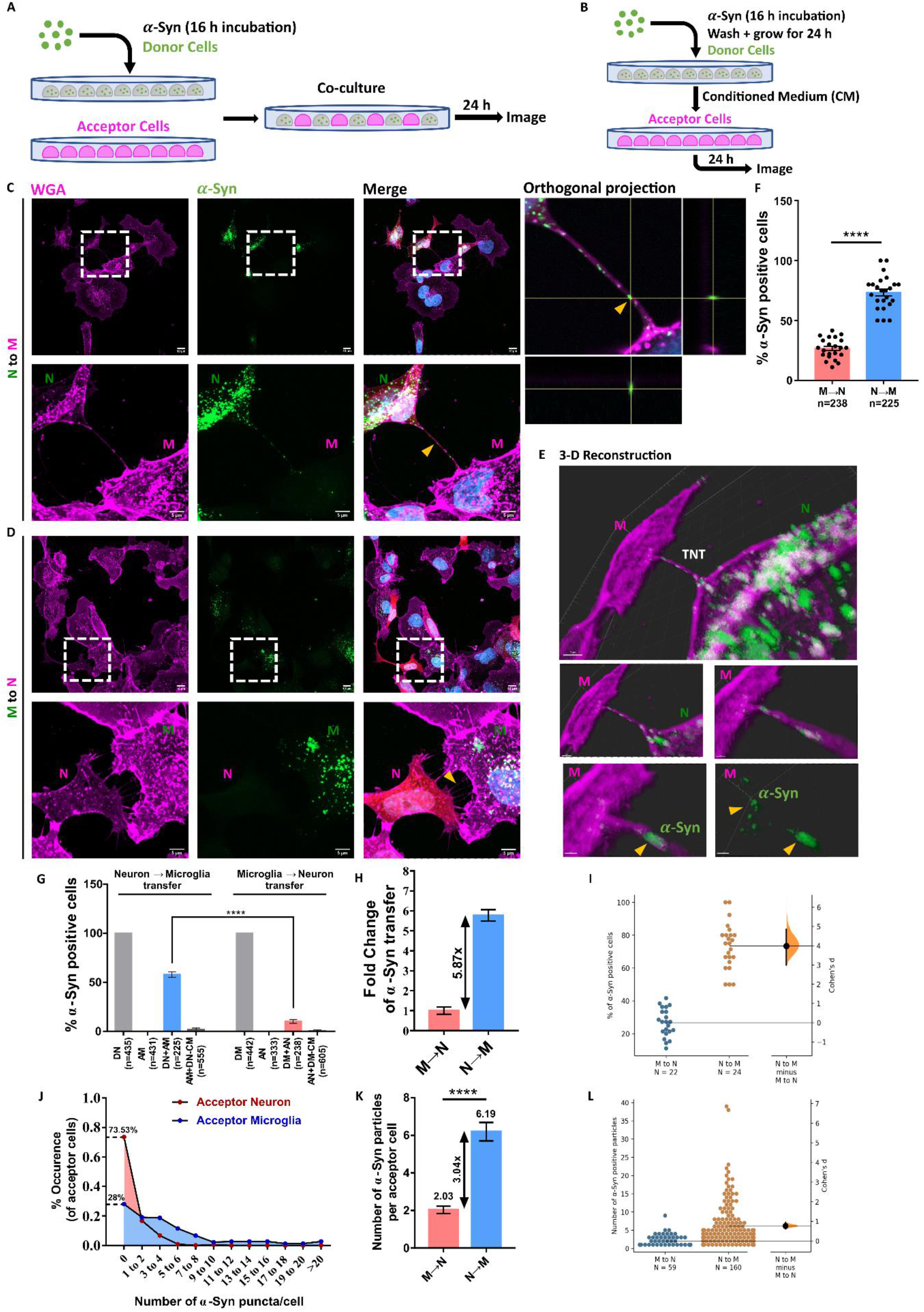
Movement of *α*-Syn between SH-SY5Y and HMC3 cells via TNTs. (A) Schematic representation of co-culture strategy used to assess for transfer. (B) Schematic representation of secretion control. (C) In a co-culture system, TNTs connect acceptor microglia with donor neuronal cells (N→M transfer). Lower panels are zoomed images of the ROI boxed in the upper panels. Yellow arrowhead in the orthogonal projection panel depicts *α*-Syn puncta inside TNT. (D) Co-culture of donor microglia with acceptor neuronal cells (M→N transfer) show the presence of TNTs between two cell types (lower panels), but without any *α*-Syn puncta within them. (E) 3-D reconstruction of connected cells in (C), done with Imaris software. M represents acceptor microglia and N represents donor neuronal cell. (F) Proportion of acceptor cells positive for *α*-Syn puncta in a co-culture system. N=3, n=238 cells for M→N transfer and n=225 cells for N→M transfer. ****p<0.0001, unpaired Student’s t-test. (G) Compiled representation of *α*-Syn transfers between both cell types, normalised for secretion control. ****p<0.0001, One-Way ANOVA. DN: Donor Neuronal cells, AM: Acceptor Microglia, DM: Donor Microglia, AN: Acceptor Neuronal cells, CM: Conditioned Media. (H) Fold-change difference between the two groups (M→N and N→M transfers) indicate a 5.87-times increase in the movement of *α*-Syn from neuronal cells to microglia. (I) Estimation statistics plot corresponding to (F). (J) Distribution pattern of the number of *α*-Syn puncta transfer in acceptor cells. Higher value for “0” is indicative of more instances of no transfer. (K) Average number of *α*-Syn particles per acceptor cell. N=3, n=59 cells for M→N transfer and n=160 cells for N→M transfer. ****p<0.0001, Student’s t-test. (L) Estimation statistics plot corresponding to (K).

73.53 % of acceptor neuronal cells did not receive any *α*-Syn from donor microglia, whereas only 28 % of acceptor microglia were negative for *α*-Syn signal (Fig. 4G). Importantly, these values were all normalized with secretion control (see materials and methods) (Fig. 4B, S1B). The extent of secretion-based transfer was, however, only marginally increased (by 1.23-folds) from neuronal to microglial cells (Fig. 4J), supporting the relevance of cell to cell mediated pathway for N→M transfer aggregate transfer over secretion. To directly demonstrate that TNTs were effectively transferring *α*-Syn aggregates between cocultured cells we used live confocal microscopy. By this approach we could observe *α*-Syn puncta moving in TNTs from neuronal cells to microglia over a period of 30 minutes (Fig. S2, supplementary movie 1). Overall these data suggest that burdened neuronal cells actively relay *α*-Syn aggregates to microglia mainly using TNT-mediated transfer The contrary, however, is not true, suggesting that microglia might play a neuroprotective role by accepting these aggregates, rather than contributing to aggregate spreading.

### Microglia provide mitochondria to neuronal cells

From our observation of a skewed directionality of *α*-Syn transfer, we next asked the possible functional significance of TNTs from a microglial perspective, and whether microglia could use TNTs to preferentially transfer materials to SH-SY5Y cells. An important pathological consequence of *α*-Syn accumulation in neuronal cells is mitochondrial damage. With the hypothesis of microglia providing neuroprotective functions to neuronal cells, we tested the possibility that microglia could supplement mitochondria to *α*-Syn-loaded SH-SY5Y cells. We co-cultured HMC3 labelled with MitoTracker Red CMXRos (donor cells) and SH-SY5Y with (+*α*-Syn) or without (WT) fibrils (acceptor cells) (Fig. 5A). Microglial conditioned medium was added to both acceptor neuronal cells to normalise for secretion control (Fig. 5B). As observed previously, functional TNTs were observed between neuronal cells and microglia (gray connections, pseudo-coloured), containing mitochondria (orange particles, pseudo-coloured) within them (Fig. 5C-F). We observed a 6.27-folds increase in mitochondrial transfer to *α*-Syn loaded (45.95 %) as compared to WT SH-SY5Y cells (7.33 %) (Fig. 5G, I, L). On the other hand, secretion-mediated transfer was comparable for both the groups (Fig. 5H). Besides the higher percentage of acceptor cells, we also observed a 1.68-folds increase in the number of mitochondrial particles received by *α*-Syn loaded neuronal cells compared to WT cells (Fig. 5K) (average 3.34 mitochondrial particles in *α*-Syn loaded cells compared to 1.98 particles in WT cells). Furthermore only 30.1 % of the *α*-Syn loaded cells did not receive any mitochondria, while 71.27 % of WT cells were negative for any mitochondrial signals (Fig. 5J, M). Using time-lapse imaging, could demonstrate the movement of mitochondria in TNTs from microglia to *α*-Syn-loaded neuronal cells (Fig. S3, Supplementary movie 2). Of interest, we could also observe co-transfer of *α*-Syn aggregates from neuronal cells to microglia, and mitochondria from microglia to neuronal cells along the same tube (Fig. S4, Supplementary movie 3). Taken together, bidirectional transfer of *α*-Syn and mitochondria is indicative of active distribution of materials between the connected cells, wherein unhealthy SH-SY5Y cells receive metabolic support from microglia, which in turn receive “toxic” aggregates.

**Fig. 5:**
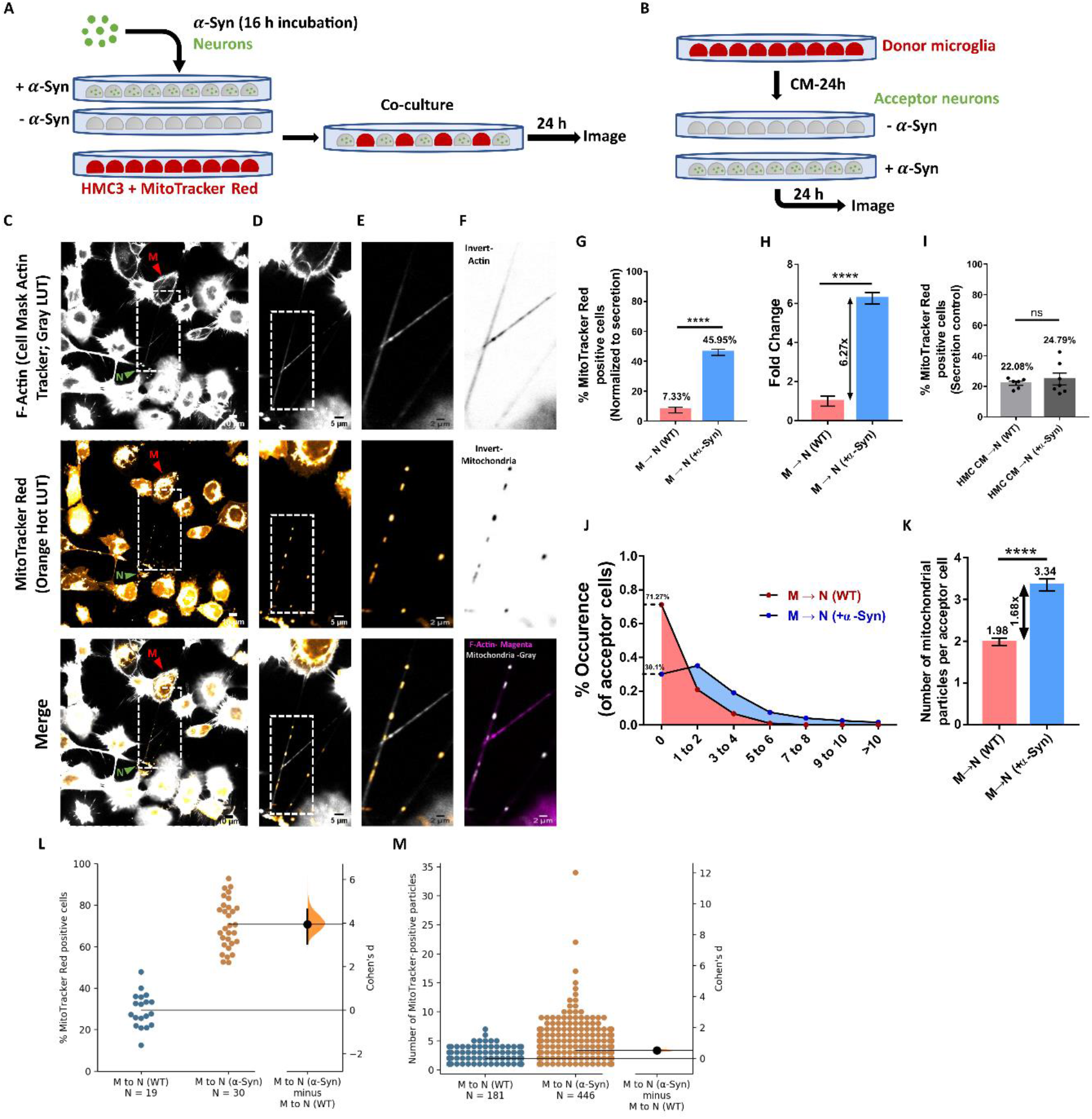
Movement of mitochondria from HMC3 to SH-SY5Y cells. (A) Schematic representation of the co-culture strategy used to assess transfer. (B) Schematic representation of secretion control. (C-F) Single, middle stack images of neuron-microglia co-culture stained for F-Actin with CellMask deep red Actin tracking stain (pseudo-coloured gray, upper panels) and mitochondria with MitoTracker Red CMXRos (pseudo-coloured orange, middle panels). White, dashed boxes indicate the ROI for zoomed images. Images were acquired using Nikon Eclipse Ti2 spinning disk microscope. (G) proportion of healthy-M→N (WT) and unhealthy-M→N (+*α*-Syn) acceptor cells positive for mitochondrial particles. N=3, n=630 healthy (WT) acceptors and n=638 unhealthy (+*α*-Syn) acceptors. ****p<0.0001, unpaired Student’s t-test. (H) Fold change of difference between the two acceptor populations in receiving mitochondrial particles from microglia depict a 6.27-times increase for unhealthy neuronal acceptor population. ****p<0.0001, unpaired Student’s t-test. (I) Proportion of healthy-M→N (WT) and unhealthy-M→N (+*α*-Syn) acceptor cells positive for mitochondrial particles in secretion control. N=3, n=634 healthy (WT) acceptors and n=605 unhealthy (+*α*-Syn) acceptors. ns: non-significant, unpaired Student’s t-test. (J) Distribution pattern of the number of mitochondrial puncta transfer in acceptor cells. Higher value for “0” is indicative of more instances of no transfer. (K) Average number of mitochondrial particles in acceptor neuronal cells. N=3, n=181 for healthy (WT) acceptors and n=446 for unhealthy (+*α*-Syn) acceptors. ****p<0.0001; unpaired Student’s t-test. (L-M) Estimation statistics plots corresponding to un-normalised transfer in (G) and (J) respectively.

### α-Syn aggregates increase TNT formation

Finally, we asked whether the presence of aggregates would affect TNT formation between microglia, as shown previously for CAD neuronal cells (27). We therefore incubated HMC3 cells with *α*-Syn for 16 h previous assessing TNT formation. Compared to control conditions, (Fig. 6A-top panel), we found a 21.92 % increase in the proportion of TNT-connected microglia in the *α*-Syn treated group (Fig. 6A- bottom panel, 6B, C). Besides the percentage of connected cells, an important question that arises is if the number of connections between cells also increase upon *α*-Syn exposure. We found an increase of more than 2-folds in the number of TNTs per connected pair when cells were treated with *α*-Syn (Fig. 6D, E), suggesting that microglia alter their morphological profile to form intercellular connections upon exposure to *α*-Syn.

**Fig. 6:**
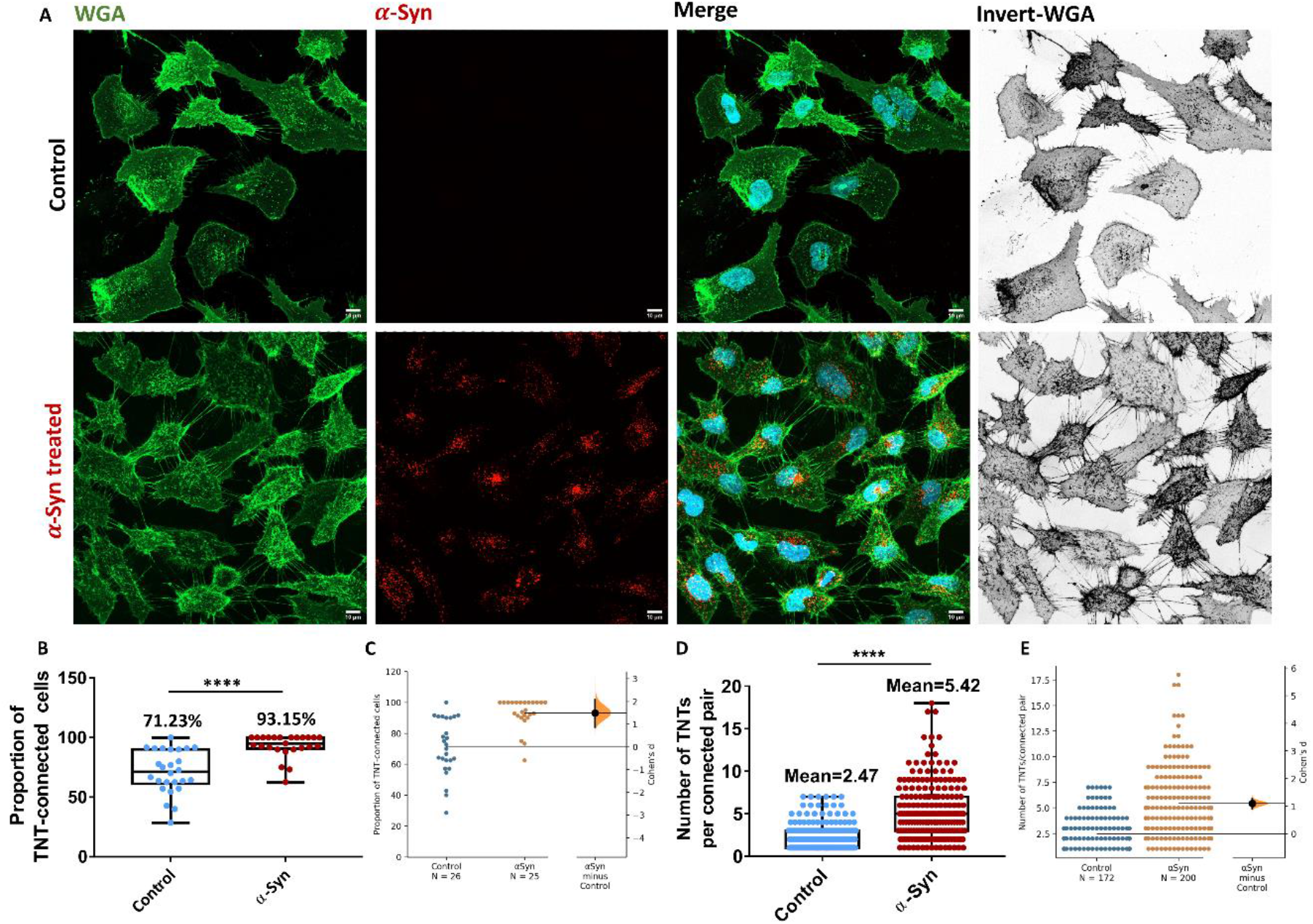
*α*-Syn exposure increases TNTs between microglia. (A) Microglia stained for membrane with AlexaFluor488-conjugated WGA (green) depicts an increased global connectivity between cells in the presence of *α*-Syn (lower panels), as compared to the control group (upper panels). (B-C) Proportion of TNT-connected cells (B) and corresponding estimation statistics plot (C). N=3, n=252 cells for control group, n=345 cells for *α*-Syn group. (D-E) The average number of TNTs between connected pairs (D) and corresponding estimation statistics plot (E). N=3, n=172 connected pairs for control group, n=200 connected pairs for *α*-Syn group. ****p<0.0001, unpaired Student’s t-test.

## Discussion

TNTs represent a fairly new mode of communication between cells of the same or different types. First reported to be formed between PC12 cells, subsequently several other cell types have been shown to form TNTs, supporting the potential involvement of these long-range membranous channels in health and diseases. TNTs have been implicated to play crucial roles early during development (42), as well as in neurodegenerative diseases, classically manifesting later in life (43). TNT-mediated transfer of protein aggregates from an unhealthy to a naïve cell aid in the propagation of the pathology by allowing the seeding of aggregates in acceptor cells (28). On a parallel tangent, TNTs can also facilitate the dilution of aggregates in a cell by sharing it with connected cells for eventual degradation. We have previously shown that neurons transfer *α*-Syn aggregates to astrocytes via TNTs where these fibrils were degraded (29). Besides astrocytes, microglia are major, yet controversial, players in NDD pathologies and their progression. On one hand microglia can engulf and degrade *α*-Syn fibrils released by neurons to prevent neurodegeneration (19), on the other hand, these cells can also directly (via release of matrix metalloproteases) and indirectly (via secretion of neurotoxic factors) influence death of neurons (6). Of interest, an elegant study has shown that microglia form TNT-like connections with each other, enabling the distribution of *α*-Syn aggregate load between microglia and their subsequent degradation (32). These data could represent a double-edged sword: if microglia were to transfer *α*-Syn aggregates to neurons, despite microglia’s ability to degrade aggregates, it would contribute to the spread of pathology. On the other hand, if there were a unidirectional transfer of aggregates from neurons to microglia, then these cells would perform a protective function. Thus, the question arises as to what the extent of TNT-mediated intercellular communication between neurons and microglia is, and the involvement of the latter in NDD pathology prevention and/or spreading. Heterotypic TNTs have been show to facilitate transfer of aggregates between different cell types (such as neurons-astrocytes and neurons-pericytes) (29,31). However, no study has assessed the presence and role of TNTs between neurons and microglia.

Our quantitative data demonstrate that around 40 % of microglia cells in culture are connected by TNTs under normal growing conditions, highlighting the potential importance of these structures (Fig. 1A-B). Most of the TNTs had lengths ranging from 10 – 20 *µ*m, with several of them extending up to 40 *µ*m (Fig. 1C, D). Despite previous reports indicating that cells of myeloid origin have thick TNTs (40,44), we found that ∼70 % of these connections were thin, F-Actin containing TNTs, while only ∼30 % were thick, F-Actin and microtubule containing TNTs (Fig. 2A-E). Furthermore the extent of *α*-Tubulin presence was categorised to be “low”, “partial”, or “complete”. Only a very low proportion of TNTs had high presence of Tubulin. The predominant presence of F-Actin is validated by the data showing that knocking out ROCK2, a regulator of Actin polymerisation, significantly increases the exchange of *α*-Syn aggregates between microglia (32). On the other hand, it will be interesting to relate the cytoskeletal composition with the formation of TNTs, as to whether the presence of tubulin polymers indicates a maturation of these structures, and/or with the transfer abilities of TNTs.

Here we show that microglial TNTs are functional as they contain mitochondrial particles and support their transfer between cells (Fig. 3A-D, supplementary movie 3). These data support recent findings showing transfer of *α*-Syn aggregates in TNTs between microglia (32). To understand whether microglia are involved in the spreading of aggregates to neurons or exert a protective function by up taking *α*-Syn aggregates from neurons, we carried quantitative and qualitative microscopy studies in co-culture of neuronal and microglial cells. We demonstrate that functional TNTs form between microglia and neuronal cells. Quantitative assessment of the transfer of *α*-Syn aggregates show that the extent of aggregate transfer is quite low from microglia to neuronal cells (Fig. 4D, F, G), while we observe a 5.87-folds increase in the transfer from neuronal to microglial cells (Fig. 4H). Given a major function of microglia is phagocytosis and degradation of pathogenic materials, we speculate that this could be a way for neuronal cells to lower their aggregate burden, suggesting that microglia could exert a protective function for neurons, similar to what we had previously proposed for astrocytes, rather than contributing to the spreading of the pathology (29). This hypothesis is supported by the recent finding that acceptor microglia are involved in active degradation of *α*-Syn aggregates after receiving it from donor cells (32). Furthermore, we confirm and extend previous findings showing that microglia increase intercellular connectivity in the presence of *α*-Syn aggregates. Not only is the global connectivity increased (Fig. 6A-lower panels, B, C), the number of TNTs connecting two cells are also increased by 2.19-folds (Fig. 6D, E). In the presence of *α*-Syn aggregates, the molecular landscape of microglia changes from a homeostatic to a reactive state, with upregulation of pro-inflammatory genes, including chemotactic genes (32). We speculate that such drive of microglia to a pro-inflammatory state might potentially alter the cellular cytoskeleton dynamics of these cells, allowing them to form more intercellular connections with each other, similar to what has been shown for HIV-1 infected macrophages (45) and dendritic cells (46). The (patho)physiological advantage of such connections being exchange of aggregates for eventual clearance. *α*-Syn-mediated increase in connections could therefore be a dose-dependent (concentration of α-Syn) or a time-dependent (period of incubation) process, both of which need to be further elucidated.

Next we asked the question whether microglia would use TNTs as donor cell to supply material to healthy or unhealthy neurons. We observed a significant increase (6.27-folds) in the transfer of mitochondrial particles from microglia to unhealthy neuronal cells, as opposed to healthy ones (Fig. 5G, 5I). Interestingly in PC12 neuronal cells, transfer of mitochondria from a healthy cell to an unhealthy, UV-irradiated cell has been reported to rescue the phenotype of the latter (47). Additionally, transfer of functional mitochondria from healthy pericytes to oxygen and glucose deprived (ischaemic) astrocytes via TNTs rescue the latter from apoptosis (48). Thus, we speculate that this could be a possible way of rescuing metabolic health of *α*-Syn burdened cells. Interestingly, we observed simultaneous, bidirectional transfer of *α*-Syn and mitochondria (Supplementary movie 3), suggestive of active interactions between the two cells working in concert to restore homeostatic balance. Ultrastructural analysis of neuronal TNTs using correlative FIB-SEM have identified bundles of individual TNTs (iTNTs) that connect the opposing cells and are held together by N-Cadherin. Such iTNTs could be aligned in parallel or anti-parallel fashions (49). Consequently, our observation of bidirectional cargo movement can be facilitated by iTNTs that maintain opposing polarities, thereby allowing unidirectional material transfer in both directions. Alternatively this could be mediated by a single, thick TNT containing both actin and microtubules.

In conclusion, our work demonstrates for the first time the presence of functional TNTs between neurons and microglia, that allow for material transfer in a selective manner. Protein aggregates are transferred with a high degree of bias from neuronal cells to microglia, whereas mitochondria is transferred preferably in the opposite direction, thus supporting the idea of protective roles of microglia in NDDs. This paves way for further investigation using murine and human primary cells, that would give subsequent insights into the mechanisms of neurodegenerative pathology spreading, especially on the involvement of microglia in NDDs.

## Acknowledgements

The authors would like to thank all the members of the membrane traffic and pathogenesis lab for insightful discussions, Sevan Belian and Roberto Notario Manzano for critical reading of the manuscript. The authors thank Dr. Aleksandra Deczkowska, Institut Pasteur, for providing HMC3 cells and Dr Takashi Nonaka and Dr Masato Hasegawa (Dementia Research Project, Tokyo Metropolitan Institute of Medical Science, Tokyo, Japan) for providing the recombinant *α*-Syn fibrils. We also thank Reine Bouyssie, a member of the administrative staff of the Membrane Traffic and Pathogenesis Unit at Institut Pasteur for her continued support. R.C. is supported by the Pasteur-Paris University international doctoral program. This work was supported by France Parkinson (Soutien de l’Association France Parkinson 2021), Don Explore (Programme Explore de l’Institut Pasteur), ANR (ANR-20-CE13-0032-01), and FRM (FRM - EQU202103012630) To CZ.

## Availability of data and materials

All data related to the manuscript are available in the main, or supplementary figures. Data can be made available upon requests.

