## Supplementary Information for "Tunneling nanotubes between neuronal and microglial cells allow bi-directional transfer of *α*-Synuclein and mitochondria"

**Running title: Neuron-microglia communicate through tunneling nanotubes**

Ranabir Chakraborty^1,2^, Chiara Zurzolo^1,2,*^

^1^Unité de Trafic Membranaire et Pathogénèse, Institut Pasteur, Paris, France

^2^Université Paris Saclay, Paris, France

**Supplementary Figure 1**

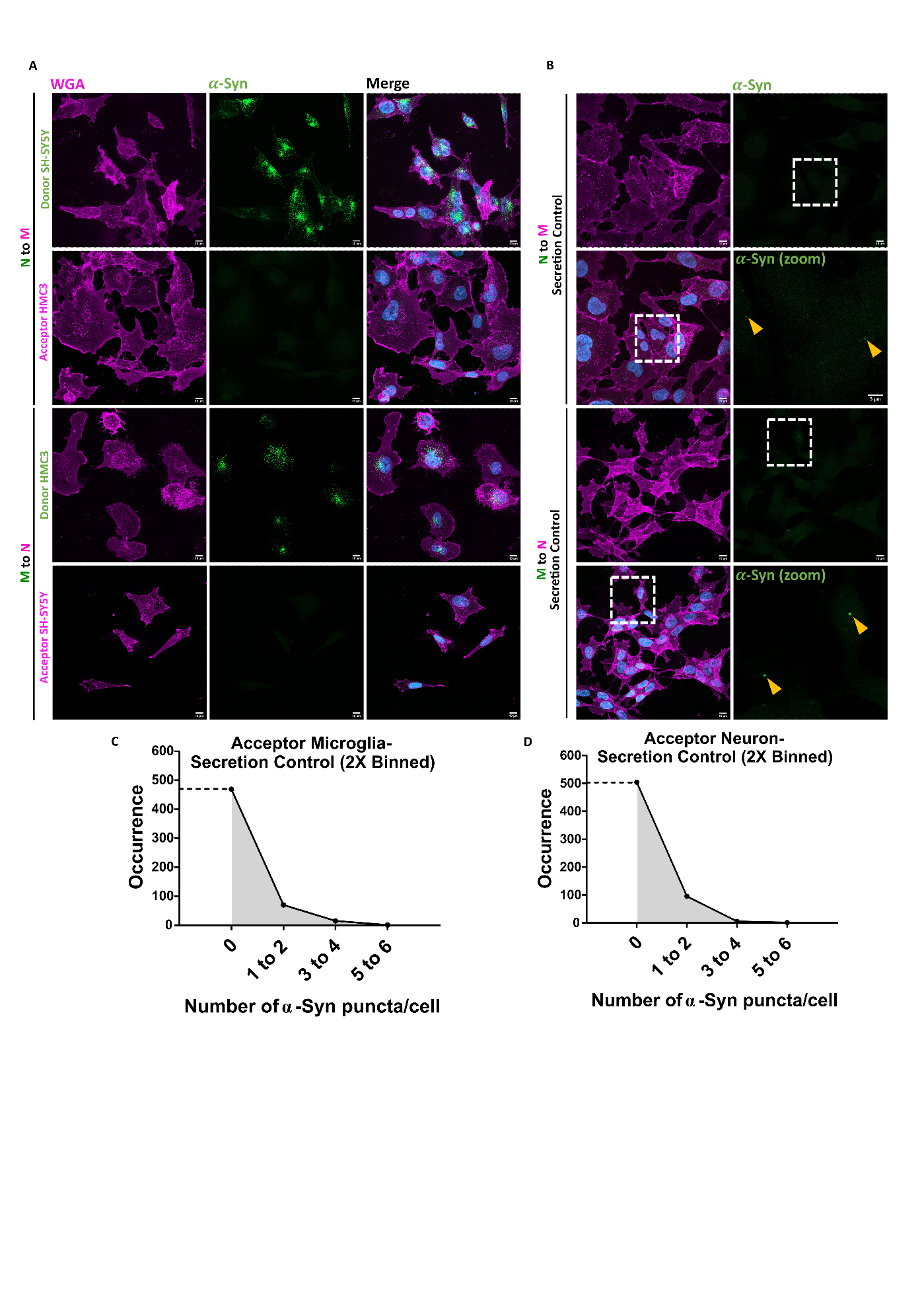

**Supplementary Fig. 1: Mono-culture and secretion control for** $\boldsymbol{\alpha}$**-Syn transfer assay between SH-SY5Y and HMC3.** (A) Mono-cultures of donor neuronal cells and acceptor microglia (upper panels, N→M transfer), and donor microglia and acceptor neuronal cells (lower panels, M→N transfer). (B) Images of acceptor cells (microglia in upper panels, neuronal in lower panels) to assess for secretion-based transfer. (D-E) Distribution pattern of the number of $\alpha$-Syn puncta in acceptor microglia (D) and acceptor neuronal cells (E) for secretion control experiments. Related to Fig. 4.

**Supplementary Figure 2**

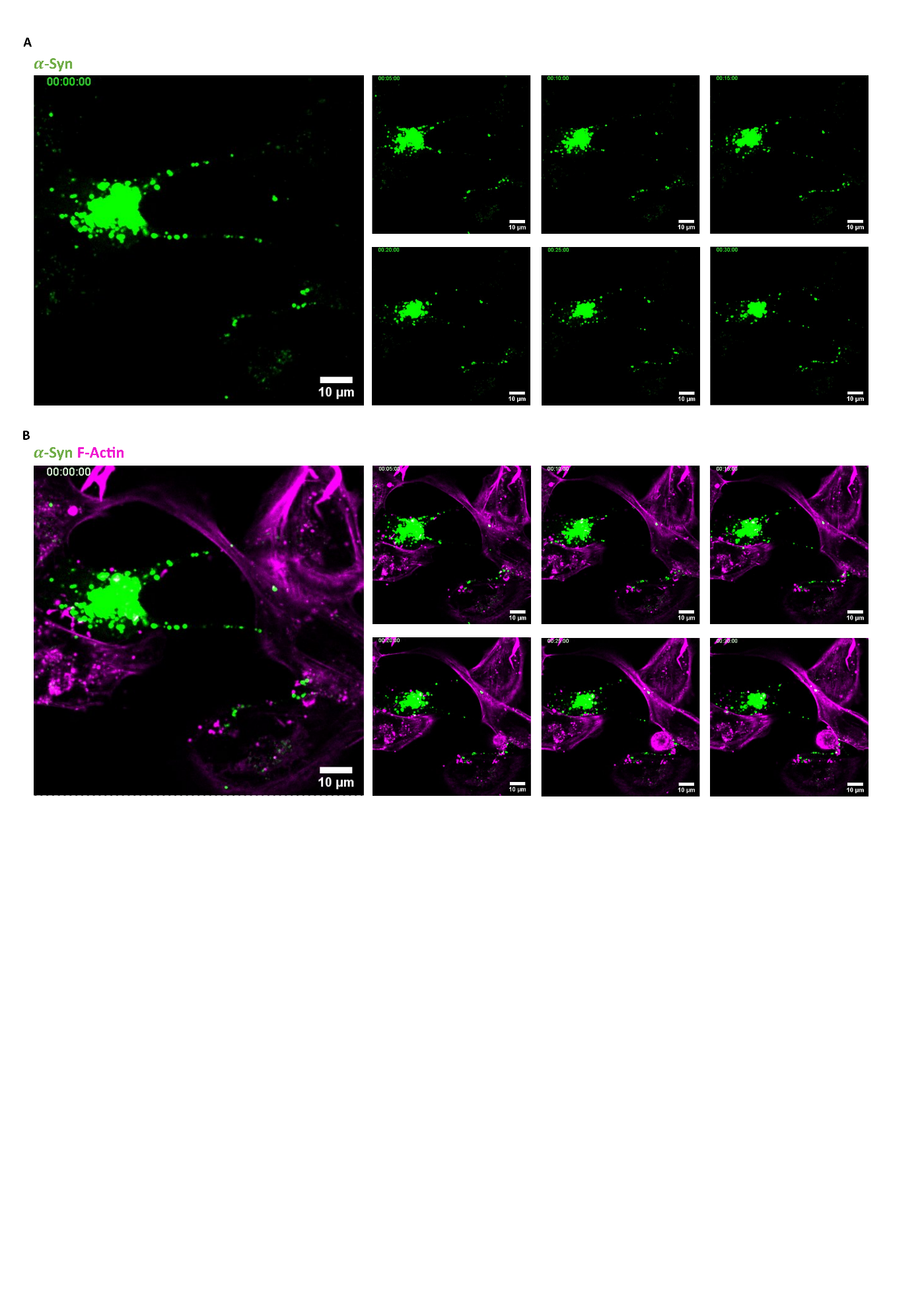

**Supplementary Fig. 2:** **Still images of movie 2.** (A-B) Time-stamps at every 5 minutes interval of $\alpha$-Syn transfer from neuronal (green $\alpha$-Syn loaded) to microglial cells. Related to Fig. 4.

**Supplementary Figure 3**

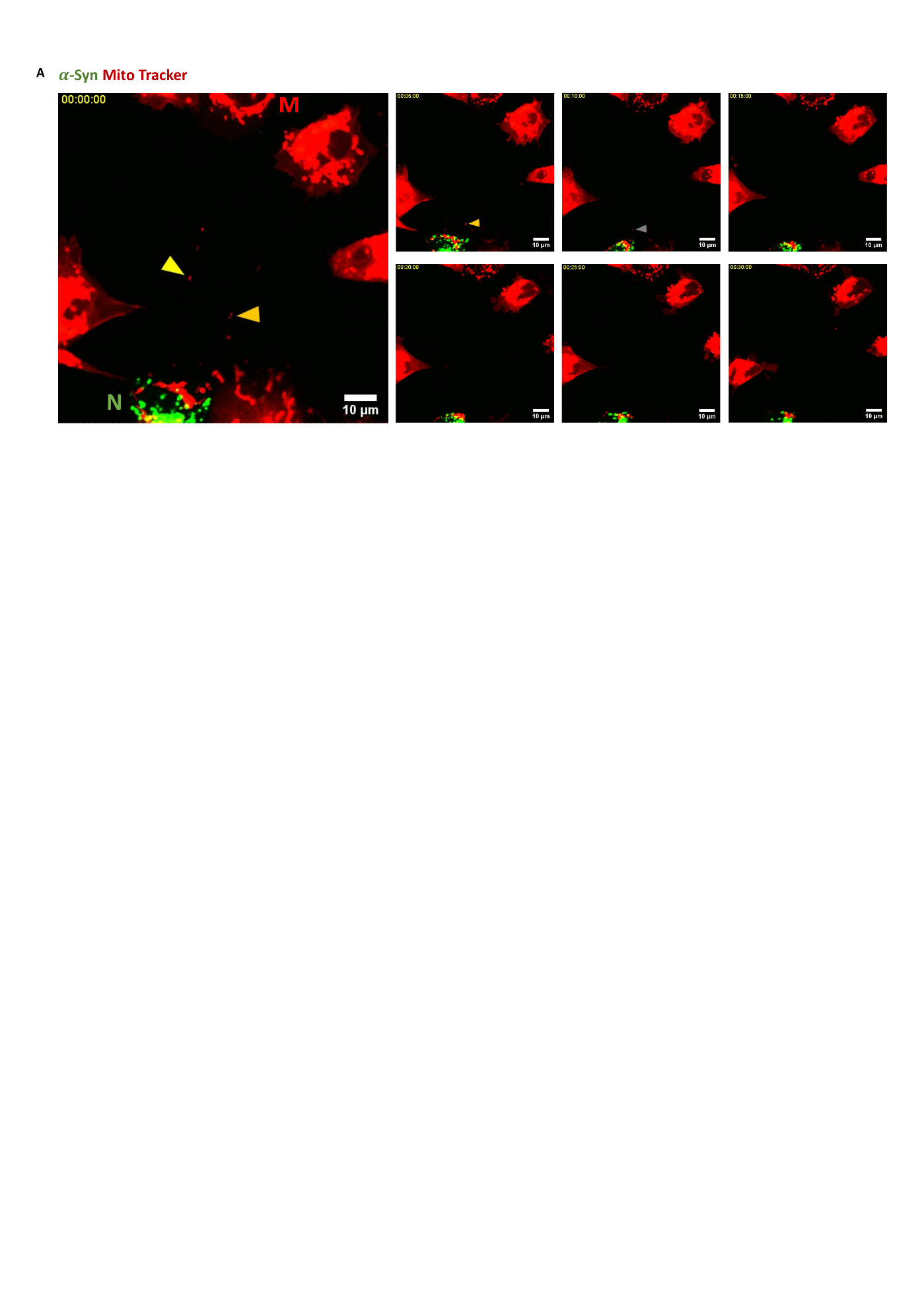

**Supplementary Fig. 3: Still images of movie 3.** (A) Time-stamps at every 5 minutes interval of mitochondrial transfer from microglia (M) to $\alpha$-Syn loaded neuronal cells (N). Related to Fig. 5.

**Supplementary Figure 4**

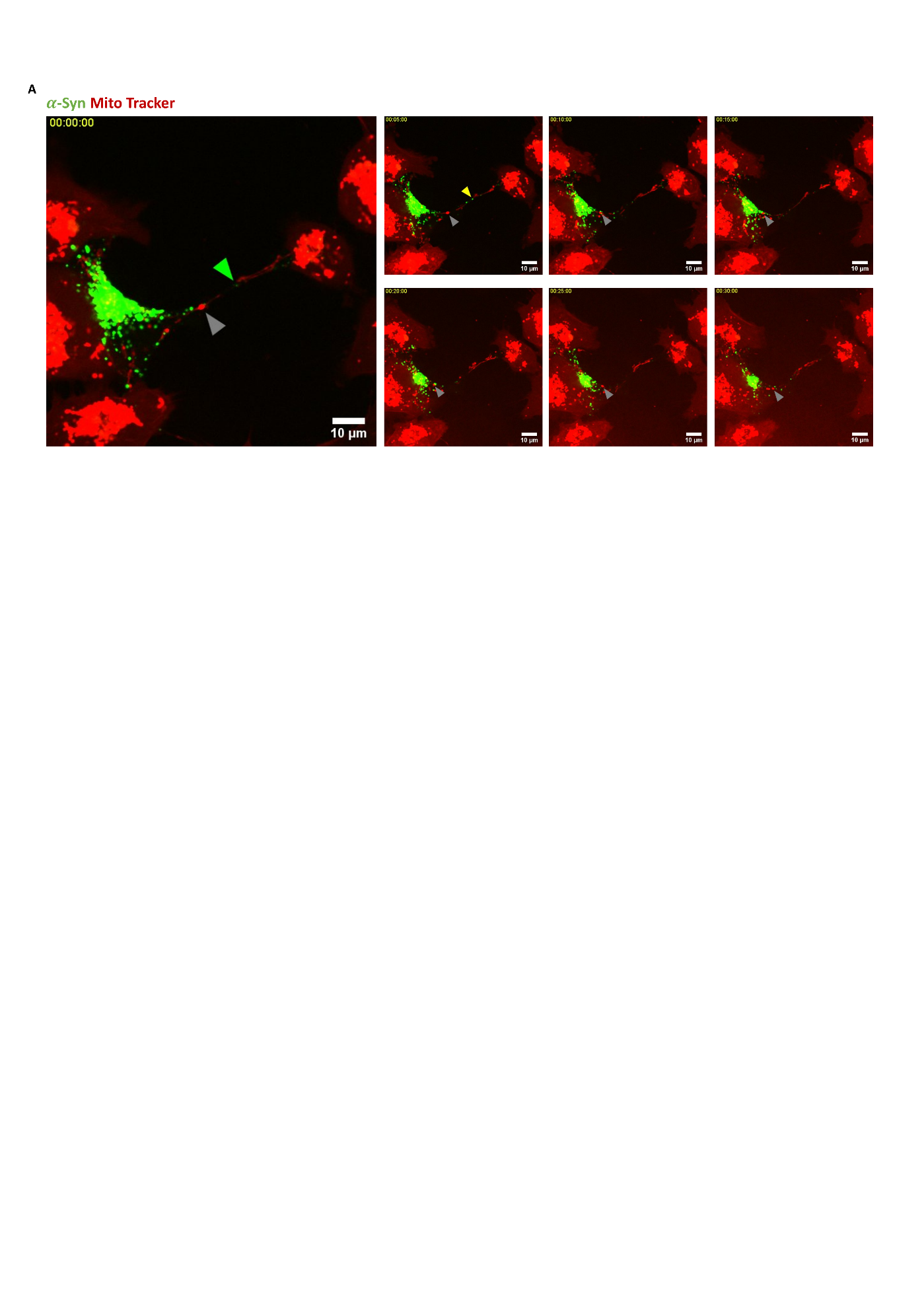

**Supplementary Fig. 4:** **Still images of movie 4.** (A) Time-stamps at every 5 minutes interval of bidirectional $\alpha$-Syn transfer from neurons (green) to microglia (red) and mitochondria from microglia (red) to neurons (green). Related to Fig. 4 and 5.

**Supplementary movie legends**

**Supplementary movie 1:** 3-D reconstruction (Imaris) of $\alpha$-Syn in TNT connecting donor neuronal cells with acceptor microglia.

**Supplementary movie 2:** Movement of $\alpha$-Syn from neuronal cell to microglia.

**Supplementary movie 3:** Movement of mitochondria from microglia to $\alpha$-Syn loaded neuronal cell.

**Supplementary movie 4:** Bi-directional movement of $\alpha$-Syn from neuronal cell to microglia and mitochondria in the opposite direction.
